# Fish predation induces drifting and emergence in an experimental stream mesocosm system

**DOI:** 10.1101/2024.07.04.602120

**Authors:** Anna-Maria Vermiert, Iris Madge Pimentel, Philipp M. Rehsen, Jonathan Meisner, Martin Horstmann, Arne J. Beermann, Florian Leese, Linda C. Weiss, Ralph Tollrian

## Abstract

Predator-prey interactions are important drivers of adaptation in aquatic communities, shaping the behaviour of invertebrates with cascading effects on community dynamics. Behavioural responses, such as moving with the downstream current (drift) or altering the timing of emergence, can be initiated and reduces the risk of encountering predators. This study aimed to examine the effects of indirect and direct predation pressure on drift behaviour and emergence patterns of aquatic invertebrates. Using an experimental mesocosm setup (*ExStream*), we analysed how the number of drifting and emerging invertebrates changed with elevated levels of chemical cues from fish (indirect predation, *Gasterosteus aculeatus* and *Cottus rhenanus*) and subsequent additional direct predation by *Cottus rhenanus* over a 9-day period. Our findings show that most of the analysed invertebrate groups displayed noticeable responses to either indirect, direct, or both forms of predation. Furthermore, our study revealed a significant impact of predation on emergence patterns, reducing the number and the size of emergent invertebrates. Given the importance of drift and emergence in dispersion and in facilitating resource flows into terrestrial systems, our findings indicate a strong effect of predation on the communities.

## Introduction

The interplay between predators and their prey in freshwater ecosystems constitutes a pivotal facet of ecological balance and community structure. Stream predators exert selective pressures on prey populations, influencing behaviours, abundance, and life-history traits. Prey organisms can perceive predators during direct predation events—such as pursuit, capture, handling, and ingestion, and indirectly through chemical cues (Åbjörnsson et al., 1997; Chivers et al., 1996; Sih, 1986; R. A. Stein and Magnuson, 1976; Weiss, 2019). Some prey organisms evolved sensitivity for chemical cues related to predation. These chemical signals facilitate early predator detection and are crucial for the induction of inducible defences and survival (Gall and Brodie, 2009; Pijanowska et al., 2020; Weiss, 2019; Weiss and Tollrian, 2018). Upon perceiving these cues, prey can develop defence strategies, with inducible defences being particularly adaptable to the threat’s level and duration (Åbjörnsson et al., 1997; Gyssels and Stoks, 2005; Helfman, 1989). The threat assessment may depend on multiple factors, including the species, diversity and co-occurrence of predators, habitat variations, diurnal cycles, as well as prey density, diversity, species and size (Andersson et al., 1986; Dahl, 1998a; Dahl and Greenberg, 1996; Douglas et al., 1994; Tollrian et al., 2015; Turner et al., 1999; Walton et al., 1977; Winkelmann et al., 2011), leading to different defence responses and their duration (Culp et al., 1991; Kasumyan, 2022; Peckarsky and McIntosh, 1998; Turner et al., 1999).

Among these strategies, behavioural changes may be most efficient as they can be quickly initiated compared to morphological alterations (Hammill et al., 2010). Behavioural adaptations may include minimizing activity to avoid detection, staying away from risky areas, seeking shelter, or swiftly fleeing upon sensing an imminent attack. In stream habitats, one such behavioural adaptation is drifting. The organisms actively release their hold to passively move with the water currents (Brittain and Eikeland, 1988; Schulz and Dabrowski, 2001). When not triggered by predators, drifting serves as a natural means of dispersal and colonization for aquatic organisms, allowing them to explore new habitats and locate resources (Brittain and Eikeland, 1988). Multiple studies have shown that the predator species may play an important role in predator-induced drifting behaviour. For example, McIntosh and Peckarsky (1999) found out that mayfly (*Baetis bicaudatus*) reduced their drift due to the brook trout (*Salvelinus fontinalis*) but increased it in presence of the predatory stonefly (*Megarcys signata*). The decision to drift may also be further influenced by the prey size with prey of bigger size drifting only at night reducing their visibility. In comparison, prey of smaller size seems less constrained, as they are less preferred by the fish predators (Allan, 1978).

Additionally, predators can alter the timing of metamorphosis in aquatic insects, although this process cannot occur abruptly, unlike drift. Aquatic invertebrates may undergo earlier emergence, allowing them to escape from the aquatic habitat and seek terrestrial refuge. The reduced time to develop and forage however often also results in smaller adult sizes (Higginson and Ruxton, 2009; Ludwig and Rowe, 1990). This strategy is well-described in the house mosquito, *Culex pipiens*, where early metamorphosis serves as the main defensive strategy against fish predation (Silberbush et al., 2019, 2015). The mosquito’s short developmental period allows it to minimize its exposure to predation by a swift transition through vulnerable life stages, thereby enhancing its chances of survival. However, this strategy seems to depend on the predator species, as Beketov and Liess (2007) discovered that the predatory bug (*Notonecta glauca*) in contrast delayed development in *Culex pipiens*. Overall, induced accelerated metamorphosis is relatively rare. In many cases, predation during the aquatic immature stages instead delays metamorphosis (Beketov and Liess, 2007; Benard, 2004; van Uitregt et al., 2012). This delay may result from the costs inherent in most defensive strategies, stemming from a trade-off between risk avoidance and other life history traits. While these strategies may enhance immediate survival against predators, they often lead to reduced food intake, thereby slowing down growth rates (Abrams and Rowe, 1996; McIntosh and Peckarsky, 2004; Sheriff et al., 2020; R. Stein and Magnuson, 1976).

So far, many studies have examined the effects of predation on drift and emergence at the level of individual prey taxa (Kasumyan, 2022) or at the scale of entire benthic communities (Dahl, 1998b; Khamis et al., 2015; Winkelmann et al., 2011). However, it is not known how individual taxa of a complete community react to increased predation risk.

We here therefore investigated the effects of indirect and direct predation on an invertebrate community within an outdoor stream mesocosm system. We quantified drift and emergence in response to fish-cue enriched water and direct predator presence over nine days. We tested four hypotheses: (1) Indirect and direct predation increases the number of drifting prey taxa. (2) The duration of predator-induced drifting behaviour is limited. (3) Predation influences the size of drifting invertebrates. (4) Predation impacts the emergence of invertebrates, affecting both the number and size of hatching individuals.

## Material and methods

### Study site

The Boye catchment, located in North Rhine-Westphalia, Germany, encompasses an area of 77 square kilometers within the Ruhr region. The Boye river is defined as a lowland river with a carbonate-rich composition and a predominant sand substrate (www.elwasweb.nrw.de; OFWK ID: DE_NRW_27726, last accessed June 20, 2024). Electrofishing activities, overseen by Dr C. Edler (Bezirksregierung Düsseldorf, Dezernat 51: Nature and Landscape Protection, Fishery), revealed a considerable abundance of sticklebacks (Gasterosteidae), a limited presence of bullheads (Cottidae), and sporadic instances of other fish species, such as bleak and sunbleak (Cyprinidae), within the study area.

### Experimental design

Established on March 4, 2022, the outdoor mesocosm system (*ExStream* system; Piggott et al., 2015) was positioned adjacent to the Boye river (An der Boy, Gladbeck; coordinates: N 51.5533°, E 6.9485°). The *ExStream* system enables precise manipulation of experimental parameters, ensuring a high level of ecological fidelity by replicating identical physico-chemical conditions to those of the Boye. The system design also allows for the migration of invertebrates, contributing to its ability to provide accurate ecological representation (Beermann et al., 2018; Piggott et al., 2015; Wagenhoff et al., 2012).

The system consisted of 16 identical flow-through mesocosm channels and was based on a scaffolding with an upper and lower level (Fig. 1). One pump (Pedrollo NGAm 1A-Pro) transported the Boye water to the system. The pumps’ intake hoses were encased in protective covers with 5 mm diameter holes to shield the pumps from larger debris and prevent fish from entering the system. The water was pumped up to the upper level where two sets of interconnected collection tanks were positioned (Fig. 1 A, B). Each set was composed of three 203 L tanks arranged in series (Fig. 1 B). Water filled these tanks, with the first two acting as sediment traps to prevent system clogging, and the last tank (header tank) distributing the water via hoses to 8 mesocosms at the lower level. The water circulated within the mesocosms, exiting through a central opening, with a flow rate maintained at approximately 2.1 L/min. The discharge from the system was directed to a retention basin.

**Fig. 1:**
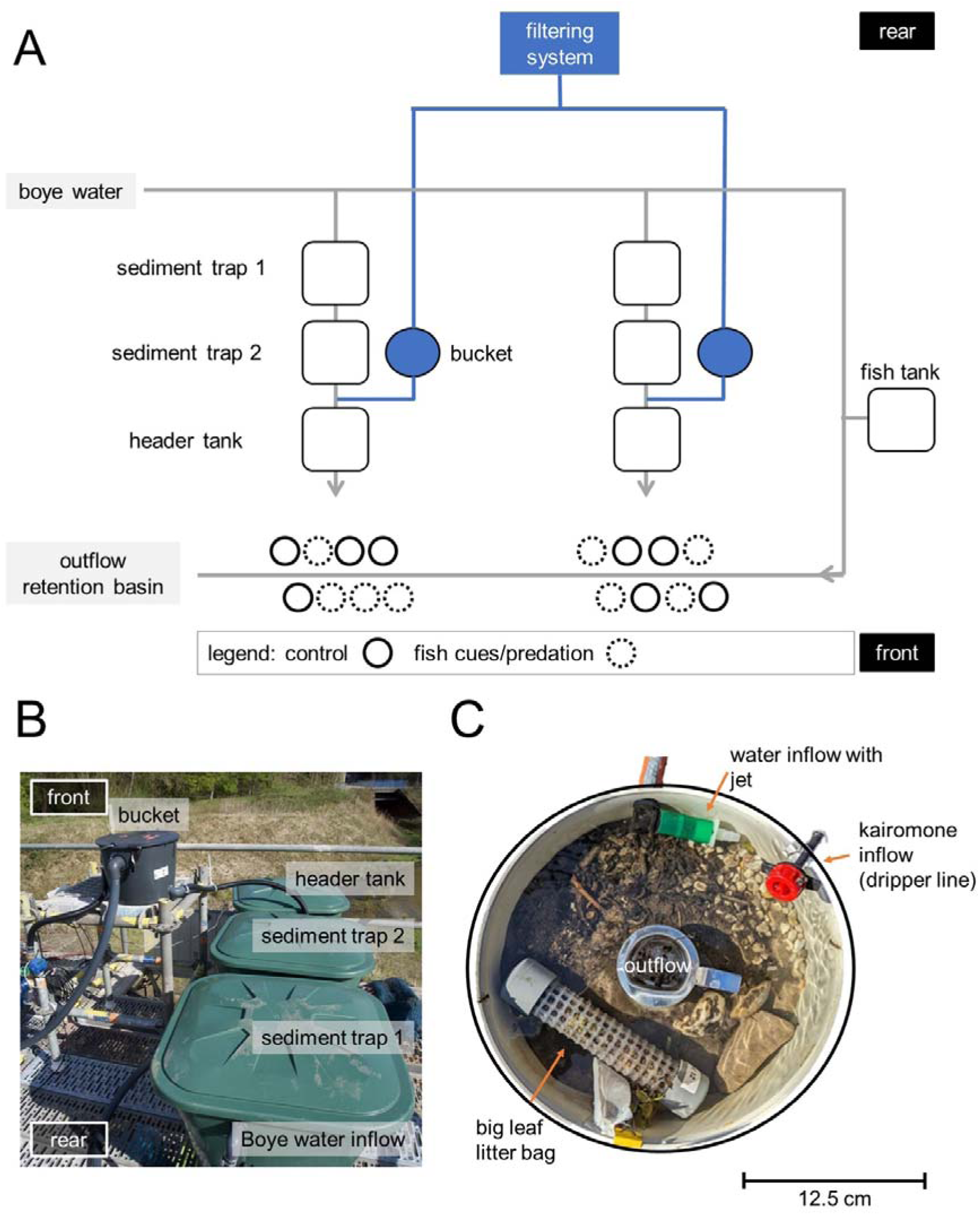
Experimental setup of the *ExStream* system (A) Schematic overview of the experimental mesocosm system with 16 mesocosms assigned in two randomized blocks. Predator-treated mesocosms are indicated by dotted circles. The direction of water flow is indicated by grey arrows. Fish-enriched water from fish tanks was applied via an irrigation line to the mesocosms. (B) Photo of the sediment traps, the header tank and the bucket at the upper level. (C) Circular mesocosm (volume 3.5 L; Microwave Ring Moulds, Interworld, Auckland, New Zealand) was filled with streambed substrate and two leaf litter bags. Water in the mesocosm had a clockwise flow direction and drained through the central outflow.

To replicate the substrates of the Boye streambed, the mesocosms were filled with a composition comprising 1000 mL of sediment obtained from the Boye catchment (N 51.5544°, E 6.9463°; sieved through a 1 mm sieve), 100 mL of fine particulate organic matter from the Boye (collected at N 51.5627°, E 6.9154°), 200 mL of quartz materials (6-8 mm), and three larger stones (40-80 mm). In addition to the sediment, each mesocosm received approximately 7 g of alder leaf litter (*Alnus glutinosa*) in a single coarse-meshed leaf litter tube (dimensions: diameter 2.5 cm, length 15 cm, mesh size 5 mm; Fig. 1 C). These tubes served as both shelter and a supplementary source of food for invertebrates. The leaves were collected during the preceding autumn and air-dried at room temperature.

### Predator exposure implementation

This study adhered to the guidelines outlined in the Animal Welfare Act. Three-spined sticklebacks (Gasterosteidae: *Gasterosteus aculeatus*) and bullheads (Cottidae: *Cottus rhenanus*), were captured and evenly distributed between two tanks (internal dimensions 113 cm x 93 cm x 57.5 cm). For each fish tank, an additional tank served as a water reservoir to maintain the water level using a mechanical pump. Sticklebacks, ranging in size from 4 to 7 cm, were captured through electrofishing at the study site. Bullheads, with a size range of 3.8 to 7.2 cm (mean: 5.72, SD: 0.78), were obtained through night fishing in the Wannebach (N 51.4245°, E 8.0511°), a tributary of the Ruhr River situated in the urban area of Arnsberg. To simulate natural feeding behaviour and natural environmental conditions, the fish in the tanks were provided a daily portion of invertebrates sourced from a downstream section.

During predator exposure implementation, invertebrates in the mesocosms were first exposed to fish-enriched water (indirect predation) and subsequently to the direct presence of a bullhead (direct predation). For indirect predation, the water from the fish tanks with an estimated total fish biomass of 480 g was used. Using a peristaltic pump (Hei-FLOW Value 01 with a multi-channel pump head C8) the water from each tank was pumped through two Tygon hoses (standard R3603, inner diameter 6.4 mm, wall thickness 1.6 mm, outer diameter 9.6 mm) into one irrigation line connected to a pressure-compensated dripper system (4 L/h per mesocosm). Diluted by the Boye water flowing through the irrigation line and the mesocosms, each mesocosm channel had a final fish concentration of 0.005 g/L.

Additionally, direct predator exposure was implemented on the fifth day of the stressor phase using bullheads (*Cottus rhenanus*). Eight bullheads were collected from the fish tanks and introduced individually into predator-treated mesocosms, thereby exposing the invertebrates to both indirect and direct predator effects. Bullheads, being benthivores, were not expected to directly influence drifting invertebrate numbers through consumption (Dahl, 1998a). This direct introduction of fish significantly increased fish biomass in the mesocosms, reaching approximately 16 mg/L, and served to increase fish cues and additionally provide visual or other cues associated with predation risk. Unexpectedly, three bullheads escaped out of our mesocosms during the experiment. It is possible that the proximity to the mating season between March and April (Kottelat and Freyhof, 2007) was too close and an important aspect of their activity was searching for a mating partner. At the end of the experiment, the escaped fish were collected in an outflow basin. All fish were returned to the area where they were collected.

### Experimental procedures

The experiment comprised two consecutive phases (Fig. 2). The first phase was a 20-day colonization period aimed at establishing natural communities. It began with the introduction of substrate into the mesocosms and the initiation of water flow. On the third day, coarse-meshed leaf litter tubes were introduced, followed by the augmentation of communities on the fifth day through multi-habitat kick-net sampling from an upstream section of the Boye, capturing macroinvertebrates from approximately 13 streambed patches (Fig. 2). The macroinvertebrates captured were introduced into the experimental mesocosms using a mixing procedure according to Elbrecht et al. (2016). The captured invertebrates were collected in a tank filled with Boye water, thoroughly stirred, and 5 litres were extracted. Eight identical jars were arranged in a circle on a potter’s wheel. As the wheel rotated, the extracted liquid was evenly distributed into the jars until they were full. The contents of two jars were then added to individual mesocosms, ensuring that each received the water and macroinvertebrates from one jar at similar densities.

**Fig. 2:**
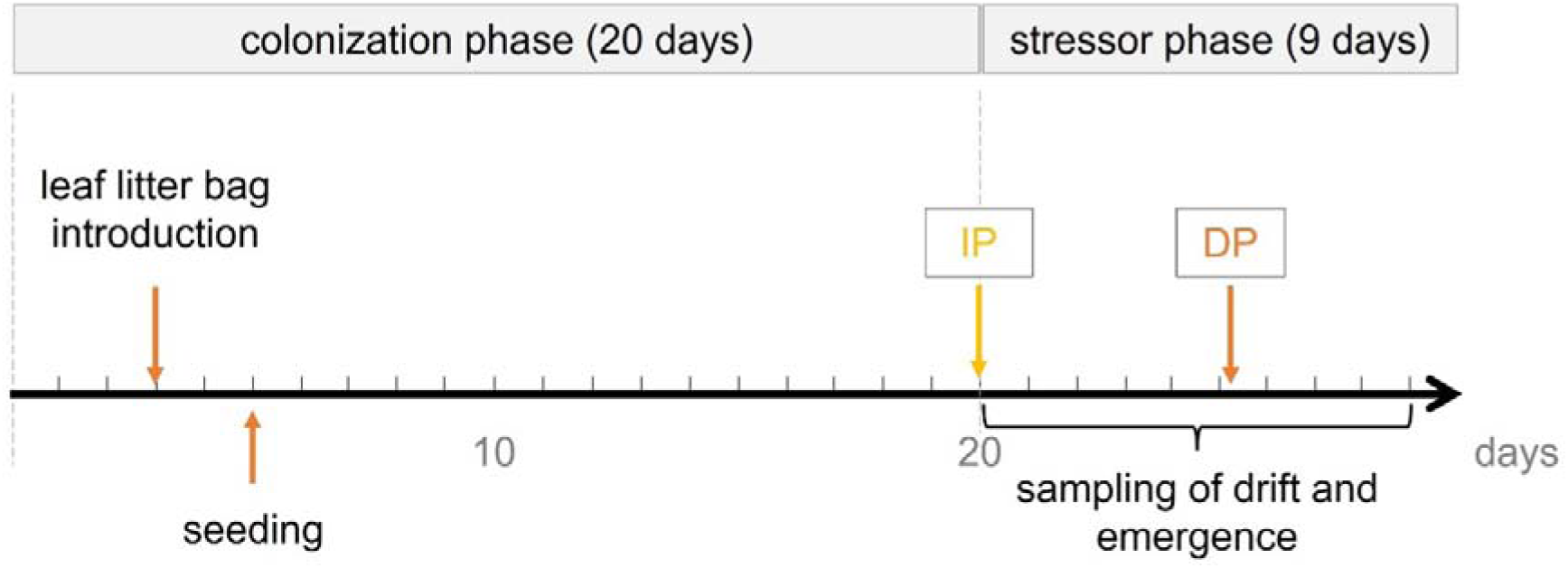
Timeline of events within the *ExStream* experiment. After setup, the 20-day colonization phase began with sediment being added to the mesocosms and water flow being activated. This was followed by leaf litter placement and a seeding event that introduced additional macroinvertebrates. The 9-day stressor phase began with the introduction of fish-enriched water (indirect predation, IP). At midnight of the fifth day, bullheads were placed in the mesocosms (direct predation, DP). The drift and emergence were continuous sampled every 24 hours at 16:00 o’clock for nine days.

Four days before the stressor phase began, 10 mm mesh nets were installed in the second sediment trap to minimize the influx of floating sediment particles reaching the header tank. Sediment was removed intermittently using a pool cleaner as needed. This process involved temporarily halting the flow within the mesocosms by closing a valve and diverting the initial water and organism influx outside the mesocosm to prevent sediment stirred up during cleaning from entering.

In the subsequent 14-day predation phase the invertebrates were exposed to first indirect predation and additionally to direct predation at the beginning of the sixth day (Fig. 2). Drift and emergence samples were collected daily for nine days starting with the stressor phase (Fig. 2).

### Sample collection and preparation

Drifting invertebrates were sampled every 24 hours and collected continuously for nine days by affixing nylon meshes to the mesocosm outflow (Fig. 3 A, B). At 16:00 each day, the drift nets were removed and preserved using 96% ethanol. In the laboratory, the invertebrates were dislodged from the drift nets under running water and collected on a 160 µm sieve.

**Fig. 3:**
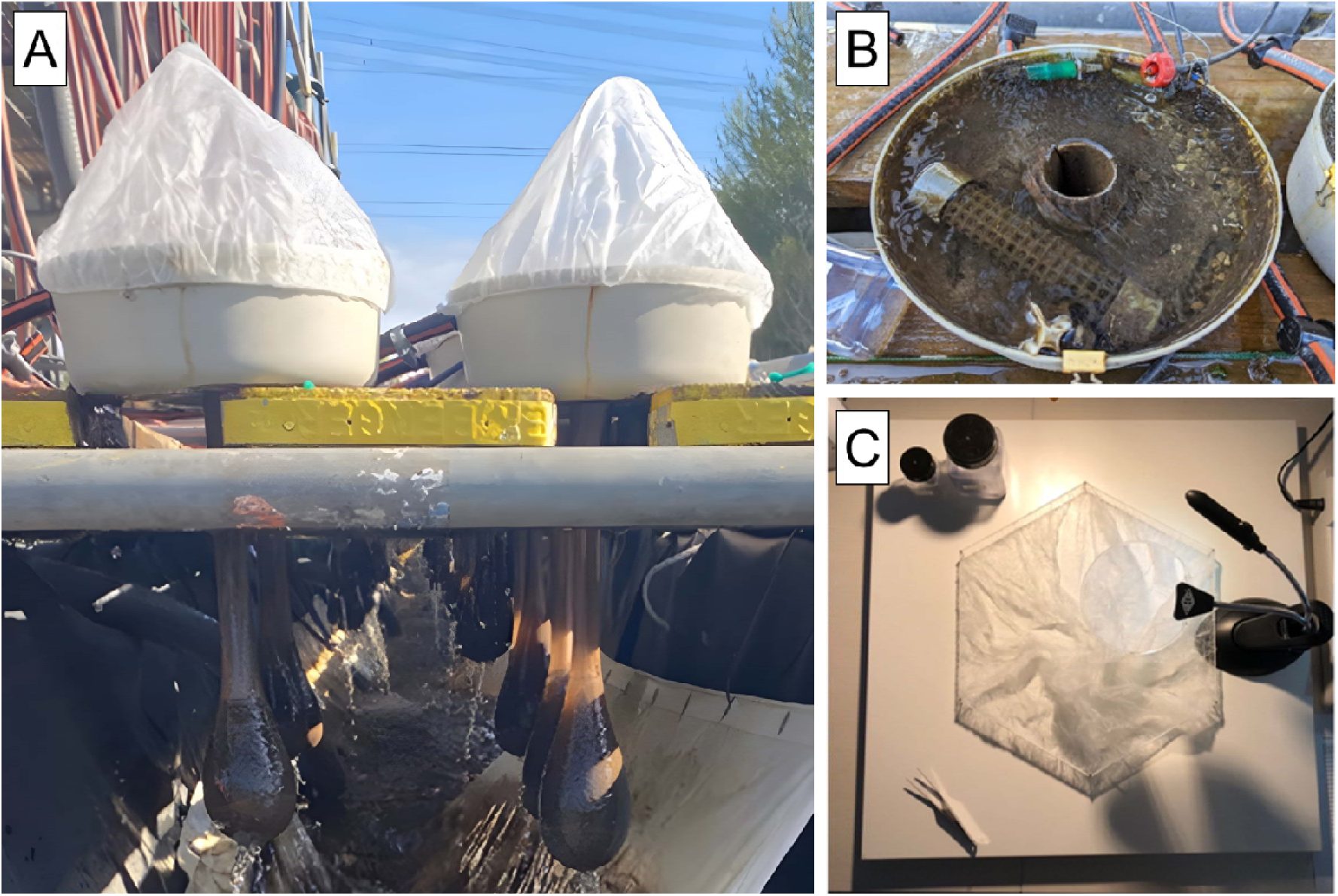
(A) Side view of the mesocosms with drift and emergence nets implemented. (B) Fixation of the drift nets on the outflow of the mesocosm in a mesocosm field experiment. (C) Nail board with affixed emergence net for easier control of imagos.

Emerging invertebrates were captured using emergence nets (Fig. 3 B) following the same schedule as drift samples and preserved similarly. These nets were installed over the mesocosms with bent wires creating an arch shape for support. In the laboratory, the emergence nets were expanded over a self-made nail board and carefully inspected for emergent insects with the aid of a magnifying glass (Fig. 3 C). This process was repeated twice to ensure meticulous inspection. The ethanol was also searched for specimens.

In both samples, invertebrates were counted and taxonomically identified based on the criteria established by Beermann et al. (2018). Invertebrates were identified to the lowest practicable taxonomic based on morphological characterisation using a microscope and standard keys (Bauernfeind and Humpesch, 2001; Eiseler, 2010; Eiseler and Hess, 2015; Lubini et al., 2012; Schaefer, 2018; Sundermann and Lohse, 2004; Waringer and Graf, 1997). To ensure statistically analysable sample sizes, rare taxa were aggregated into higher taxonomic categories. Some specimens could not be precisely identified due to their small size, leading to their remaining unidentified beyond a certain taxonomic level. The sizes of all discovered invertebrates were measured using graph paper and rounded to whole numbers. The count and size of invertebrates were used to assess the indirect and direct effects of predator exposure on drift and emergence.

### Data analysis

The statistical analysis was executed using R version 4.1.1 (R Core Team, 2021). We systematically assessed the impact of time and predator exposure on count data of both drift and emergence samples. For the drift data, the percentage composition of different drifting taxa groups was assessed over nine days under control and predation treatments, with each day’s data normalized to 100%. We analysed the changes in composition over time within each treatment and the daily differences between control and predation treatments using Fisher’s exact test. The data is presented as bar plots.

In the drift, we identified the following taxa: Crustaceans (Copepods, Anomopoda, Ostracoda, Nauplius larvae, Gammarus sp., Asellidae), Bivalves (Sphaeriidae), Gastropods (Lymnaeidae), Coleoptera (Hygrobiidae), Hirudinea (Glossiphonia sp.), larvae of the orders Diptera (Chironomidae, Simulidae, Ceratopogoninae, Stratiomyidae), Trichoptera (Hydropsychidae., Hydroptilidae., Limnephilidae, Leptoceridae), Ephemeroptera (Baetidae, Caenidae, Ephemeridae), Megaloptera, Odonata (Anispotera, Zygoptera) and Plecoptera (Nemouridae). We applied models to the total count and to distinct taxons with sufficient numbers. These taxa included Copepoda, Anomopoda, Ostracoda, Nauplius larvae, *Gammarus* sp., Chironomidae larvae, Trichoptera larvae and Ephemeroptera larvae, collectively representing over 90% of drifting individuals. Due to still low numbers for the Trichoptera, Ephemeroptera, Anompoda and Ostracoda their data was aggregated for the indirect and direct predation phases. For the total count of drifting invertebrates, organisms who were indiscernible due to either damage or insufficient size were systematically excluded from the analytical computations. Oligochaetes were also excluded from data calculations due to their dissipation into small fragments measuring less than 2 mm. For the distinct taxa analysis, the primary larval stage in crustaceans (Nauplius larvae) was collectively counted as one group due to the inherent challenges associated with accurately determining their taxa. Chironomidae pupae were excluded from the Chironomidae count due to their unlikely presence in the drift due to active drift behaviour.

The count data were fitted using the glmmTMB function from the glmmTMB package (Brooks et al., 2017). To examine drift patterns across the sampled days, we employed a Poisson regression with the identity link function (glmmTMB(abundance ∼ predation * day + (1 | Channel), family = poisson(link = “identity”), data = data)). Predation and day were treated as factorial predictors. The mesocosm channels were included as random effects to account for repeated measurements over days. Due to the change in treatment during the nine days, indirect predation (days 1 to 5) and direct predation (days 6 to 9) were modelled separately (Supplementary tab. 1, Supplementary tab. 2, Supplementary tab. 3). To address overdispersion in the models, we utilized negative binomial distributions (nbinom1, nbinom2) when necessary. The model selection was guided by comparing Akaike Information Criterion (AIC) values, choosing the distribution that provided the best fit. Model significance was assessed using Anova from the car package (Fox and Weisberg, 2019, Supplementary tab. 4). Following the detection of an interaction effect, we performed a post hoc analysis to identify the specific days where significant predation effects occurred using the emmeans package (emmeans_results <- emmeans(model, ∼ predation | day), contrast_results <- pairs(emmeans_results, adjust = “fdr”), Lenth R (2024), Supplementary tab. 5). The sample size (n = 16) remained consistent across all evaluation days, except for a missing sample on day 2 and day 9. These drift nets were lost during the experiment when they became dislodged from the outflow and were carried along by the current. Due to challenges related to the distributional properties of the Nauplius larvae count during direct predation, we added a small constant (1) to each observation (abundance_transformed = abundance + 1) to improve model fit and interpretation.

Due to the low numbers of emergences per day, it was not possible to study the emergence patterns across all nine days. Therefore, the emergence count data for days 1 to 5 (indirect predation) and days 6 to 9 (direct predation) were aggregated. We analysed the aggregated data using a Poisson regression with the identity link function (glmmTMB(abundance ∼ predation * day + (1 | Channel), family = poisson(link = “identity”), data = data)) and an Anova (Anova(model, type = II), Supplementary tab. 6). For the emergence samples, analysing different taxonomic groups was deemed impractical given the difficulties in identifying damaged specimens and the low overall number of emergences. The fitted model was also adjusted for overdispersion.

We calculated the mean body size based on rounded size measurements to investigate the potential effects of predator exposure on the size of drifting or emerging invertebrates. For drifting invertebrates, we aggregated the data by calculating the mean for days 1 to 5 (indirect predation) and days 6 to 9 (direct predation) due to the low numbers per day when separated into size classes. We tested the size of all individuals and single taxa found in the drift. Microcrustaceans were excluded from the analysis due to minimal body size differences resulting from rounding. Similarly, emergence size measurements were aggregated due to low numbers, and because changes in emergence do not occur instantaneously. This could potentially result in the effects of indirect and direct predation on size differences manifesting at later time points, leading to overlapping effects. To assess the normality of each treatment’s distribution, we used the Shapiro-Wilk test (shapiro.test()). Depending on the normality test results, we employed either a Student’s t-test (t.test()) for parametric data or a Wilcoxon rank-sum test (wilcox.test()) for non-parametric data to determine size differences between the control and predation treatments. The outcomes of the drift and emergence analyses are visually represented through boxplots, bisected at the median value, with whiskers extending to values within the 1.5 interquartile range. This graphical presentation was generated using ggplot2 from the tidyverse package (Wickham et al., 2019).

## Results

### Drift

The drift over the 9 days comprised in total 72.23% crustacean (20.63% Cyclopoida, 15.05% *Gammarus* sp., 13.85% Nauplius larvae, 9.99% Harpacticoida, 6.96% Anomopoda, 4.04% Ostracoda, 1.71 Asellidae and 0.02% others), 18.00% Diptera (9.52% Chironomidae larvae, 4.14% Tanytarsini larvae, 2.97% Chironomidae pupae, 1.37% others), 4.04% Trichoptera, 3.80% Ephemeroptera, and 1.97% others. The composition did not significantly differ over the days in either the control or predation treatment groups (Fisher’s Exact Test: control, p-value = 0.523; predation, p-value = 0.374). There was a significant difference between the control and predation treatments specifically on days eight and nine (Fisher’s Exact Test, day 8: p-value < 0.001, day 9: p-value < 0.001).

Invertebrates showed an increased propensity to drift in the mesocosms due to indirect predation, with this effect diminishing over time (LR Chi-square 14.283, df 4, Pr(>Chi-square) 0.006; Fig. 5, Supplement tab. 4). Specifically, significant differences in drift response compared to control levels were observed on the first and second days (day 1: df Inf, z-ratio 3.847, p-value < 0.001, day 2: df Inf, z-ratio 2.373, p-value 0.018; Fig. 5, Supplement tab. 5) but not in subsequent days. Direct predation also causes increased drift response, which was tendentially influenced by time (LR Chisq 7.75, df 3, Pr(>Chisq) 0.051; Fig. 5, Supplement tab. 4). Significant differences in response to direct predation were observed only on the first and second days (emmeans, day 1: df Inf, z-ratio 2.535, p-value 0.011, day 2: df Inf, z-ratio 2.016, p-value 0.044; Fig. 5, Supplement tab. 5).

**Fig 4.**
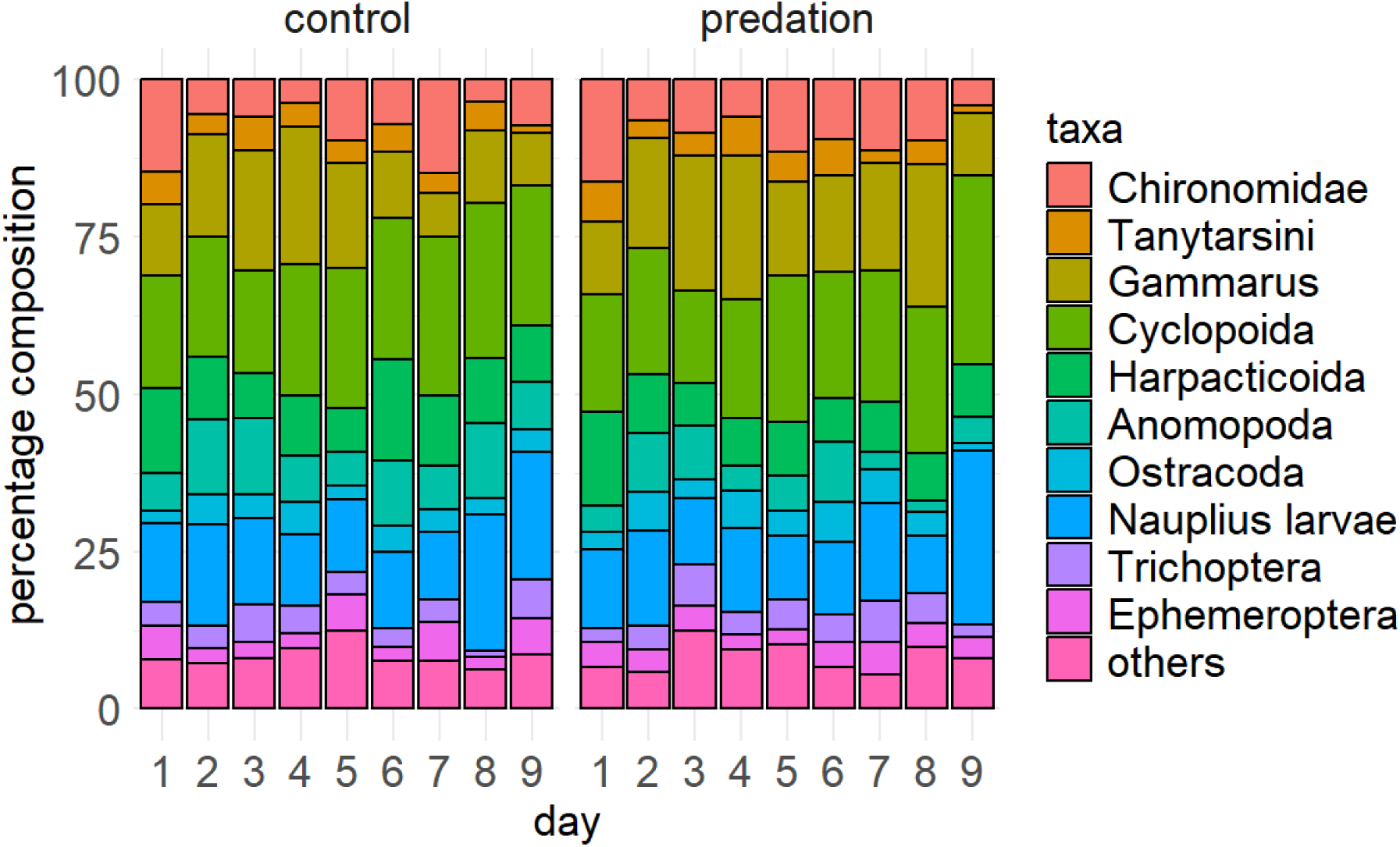
Percentage composition of different drifting taxa groups in a field mesocosm experiment. Each bar represents the relative abundance of taxa groups on a given day, normalized to 100%. Drift samples were collected over nine days for both control and predation treatments. The predation treatment was implemented by enriching the water with fish chemical cues from three-spined sticklebacks (*Gasterosteus aculeatus*) and bullheads (*Cottus rhenanus*) and introducing bullheads into the mesocosms.

**Fig. 5:**
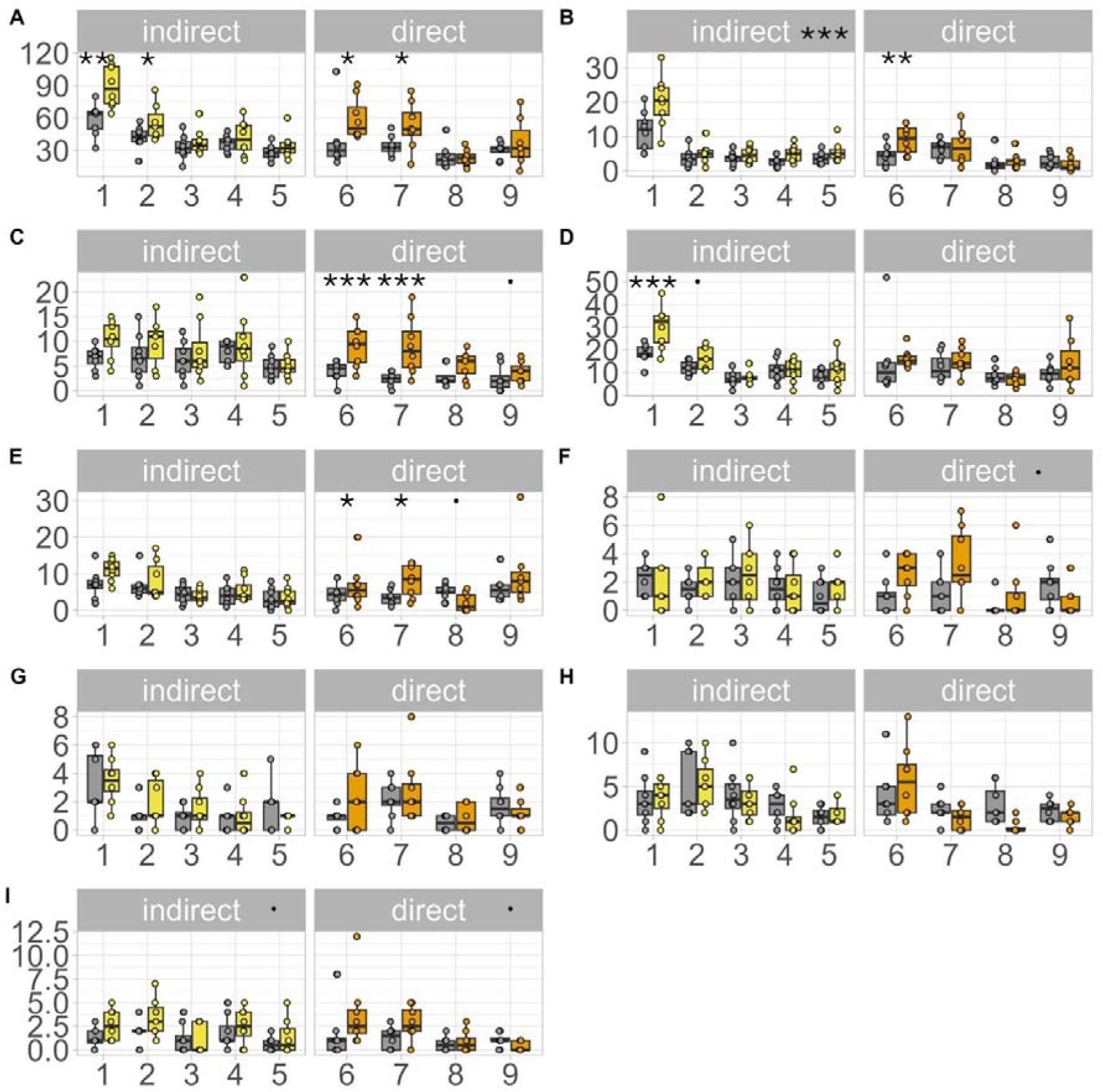
Influence of indirect and direct predation on the number of drifting invertebrates in a field mesocosm experiment over nine days compared to the control condition (grey). Water with fish chemical cues from three-spined sticklebacks (*Gasterosteus aculeatus*) and bullheads (*Cottus rhenanus*) was added continuously starting from the first day of the experiment (yellow). At midnight of the fifth day, bullheads were added as direct predation stimuli (orange). The total number of drifting invertebrates (A) and individual taxa, including Chironomidae larvae (B), *Gammarus* sp. (C), Copepoda (D), Nauplius larvae (E), Trichoptera larvae (F), Ephemeroptera larvae (G), Anomopoda (H), Ostracoda (I) were tested. Tendential and significant Anova results were noted next to the predation type. When predation and time interacted, emmeans results were displayed over the specific boxplots. Significant code: 0.1., 0.05 *, 0.01 **, < 0.001***.

Increased drift behaviour due to indirect predation was observed in some taxa. Chironomidae exhibited a significant increase in drifting across all days (LR Chi-square 10.285, df 1, Pr(>Chi-square) 0.001), while Ostracoda showed a tendency towards increased drift (LR Chi-square 3.188, df 1, Pr(>Chi-square) 0.074; Fig. 5, Supplement tab. 4). Copepods initially exhibited increased drift behaviour in response to indirect predation, but this response diminished over time (LR Chi-square 12.322, df 4, Pr(>Chi-square) 0.015; Fig. 5, Supplement tab. 4). They drifted significantly more on the first day and showed a tendency on the second day (day 1: df Inf, z.ratio 3.456, p.value 0.001, day 2: df Inf, z.ratio 1.681, p.value 0.093; Fig. 5, Supplement tab. 5). Amonopoda, Nauplius larvae, *Gammarus* sp., Trichoptera, and Ephemeroptera did not show any discernible drift response to indirect predation (Fig. 5, Supplement tab. 4).

In response to direct predation, multiple taxa exhibited increased drift behaviour. Chironomidae larvae and *Gammarus* sp. showed heightened drift responses initially, which reduced over time (Chironomidae: LR Chi-square 7.945, df 3, Pr(>Chi-square) 0.047; *Gammarus* sp.: LR Chi-square 10.533, df 3, Pr(>Chi-square) 0.015; Supplement tab. 4). Chironomidae drifted significantly more on the first day of direct predation (day 6: df Inf, z-ratio 2.663, p-value 0.008), while the drift behaviour of *Gammarus* sp. differed significantly on both the first (day 6: df Inf, z-ratio 3.339, p-value 0.001) and second days (day 7: df Inf, z-ratio 4.224, p-value < 0.0001). Nauplius larvae also exhibited changes in drift behaviour in response to predation over time (LR Chi-square 13.962, df 3, Pr(>Chi-square) 0.003). They initially responded with increased drifting on the second day (day 7: df Inf, z.ratio 2.149, p.value 0.032), significantly decreased their drift on the third day (day 8: df Inf, z.ratio −2.183, p.value 0.029), and tendentially increased drifting behaviour on the fourth day (day 9: Inf, z.ratio 1.684, p.value 0.092; Fig. 5, Supplement tab. 5). Trichoptera and Ostracoda also tendentially displayed increased drift behaviour in response to direct predation (Trichoptera: LR Chi-square 3.299, df 1, Pr(>Chi-square) 0.069; Ostracoda: LR Chi-square 3.237, df 1, Pr(>Chi-square) 0.072). Ephemeroptera, Copepoda, and Anomopoda did not show a significant response to direct predation (Supplement tab. 4).

### Size of drifting invertebrates

The microcrustaceans Copepoda, Ostracoda, Amonopoda and Nauplius larvae were the smallest individuals found, with a mean size below 1 mm. In contrast, Simulidae larvae had the largest mean size, measuring 5.3 mm (Tab. 1). The largest individuals found were a 17 mm Trichoptera and a 20 mm *Gammarus* sp. (Tab. 1).

**Tab. 1:**
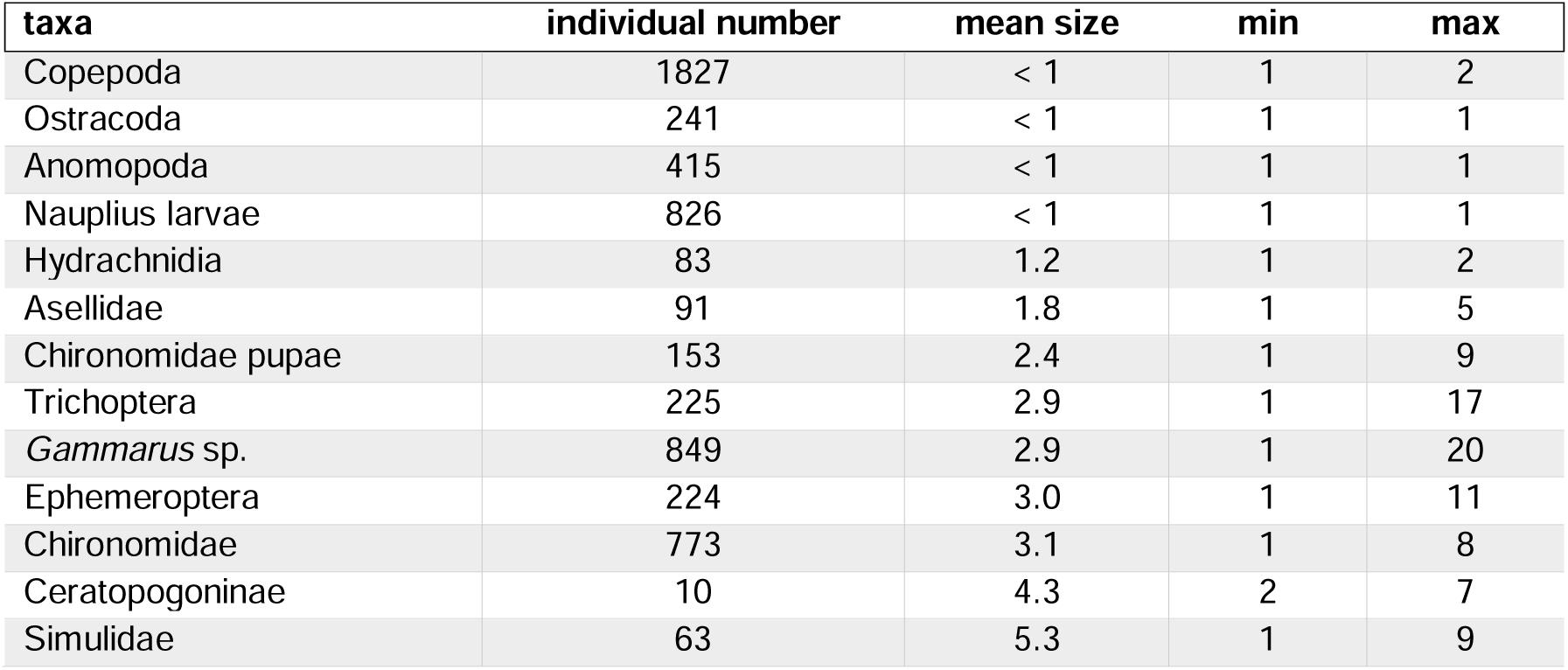
The mean, minimum, and maximum body sizes (mm) and individual counts of drifting invertebrates collected in drift during a nine-day mesocosm field experiment. Taxa with fewer than ten individuals were excluded from the analysis. Mean, minimum and maximum body sizes were calculated based on rounded size measurements.

| taxa | individual number | mean size | min | max |
| --- | --- | --- | --- | --- |
| Copepoda | 1827 | < 1 | 1 | 2 |
| Ostracoda | 241 | < 1 | 1 | 1 |
| Anomopoda | 415 | < 1 | 1 | 1 |
| Nauplius larvae | 826 | < 1 | 1 | 1 |
| Hydrachnidia | 83 | 1.2 | 1 | 2 |
| Asellidae | 91 | 1.8 | 1 | 5 |
| Chironomidae pupae | 153 | 2.4 | 1 | 9 |
| Trichoptera | 225 | 2.9 | 1 | 17 |
| *Gammarus* sp. | 849 | 2.9 | 1 | 20 |
| Ephemeroptera | 224 | 3.0 | 1 | 11 |
| Chironomidae | 773 | 3.1 | 1 | 8 |
| Ceratopogoninae | 10 | 4.3 | 2 | 7 |
| Simuliidae | 63 | 5.3 | 1 | 9 |

Indirect predation did not lead to significant size differences in the drift (t = 0.373, df = 13.139, p-value = 0.715). Larger individuals of *Gammarus* sp., Chironomidae, Trichoptera, and Ephemeroptera larvae did not increase their drift behaviour due to indirect predation (*Gammarus* sp.: t = −1.371, df = 10.619, p-value = 0.199, Chironomids larvae: t = 1.553, df = 12.167, p-value = 0.146, Trichoptera larvae: t = - 1.256, df = 11.171, p-value = 0.235, Ephemeroptera larvae: t = 0.473, df = 13.783, p-value = 0.643).

On average, larger invertebrates attempted to leave via drift due to direct predation (t = 2.830, df = 7.965, p-value = 0.022). Except for *Gammarus* sp., where larger individuals tended to drift more, larger individuals of other taxa did not show a significant increase in drifting (t = 1.916, df = 13.912, p = 0.076, Fig. 6, Tab. 2).

**Fig. 6:**
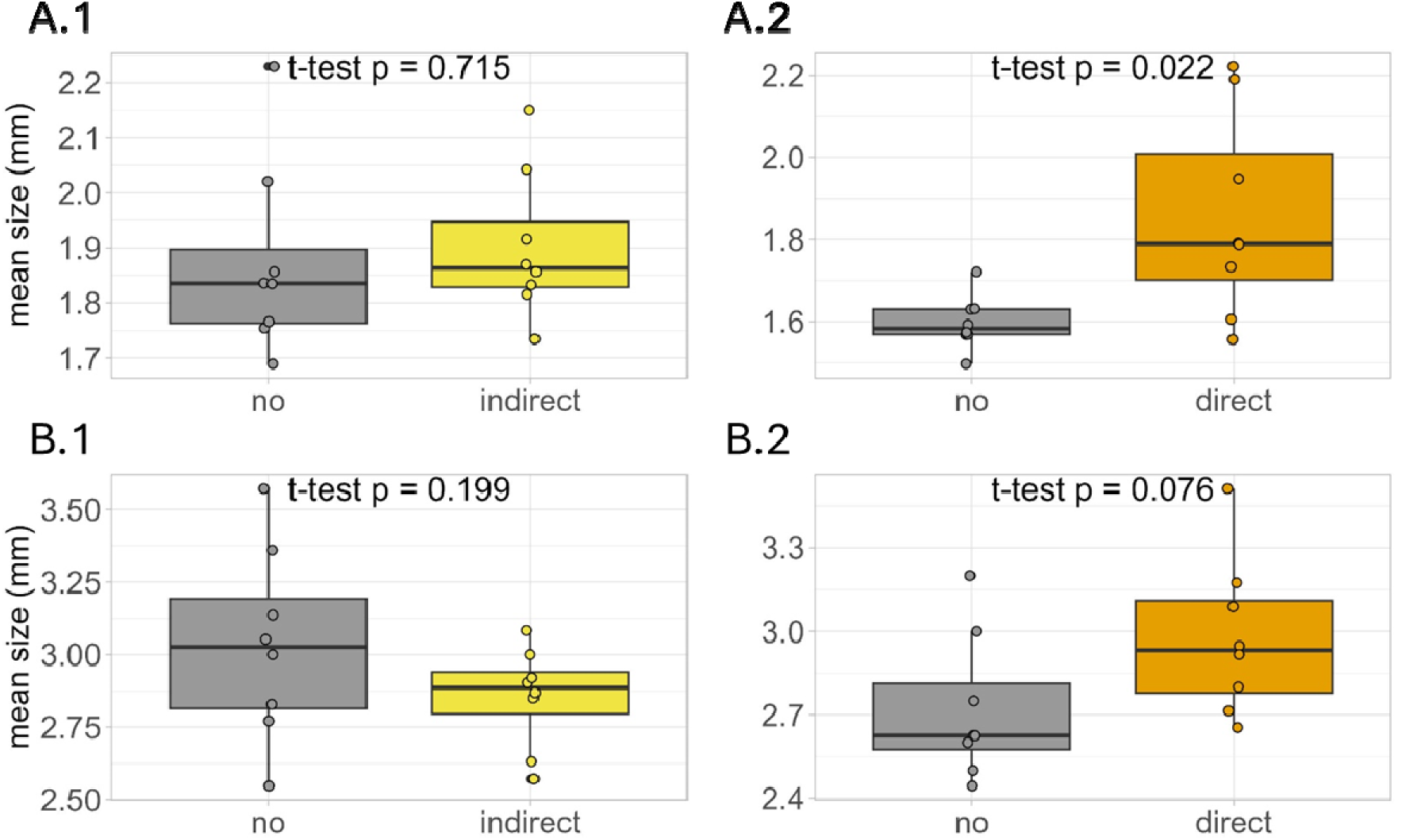
Mean size (mm) of total drifting invertebrates (A) and drifting *Gammarus* sp. (B), influenced by either indirect (.1) or direct predation (.2). We tested the drifting behaviour of invertebrates using fish-enriched water from three-spined sticklebacks (*Gasterosteus aculeatus*) and bullheads (*Cottus rhenanus*) to simulate indirect predation (yellow) and the direct presence of bullheads for direct predation (orange) in a mesocosm field experiment. These treatments were compared to a control treatment (grey).

**Tab. 2:**
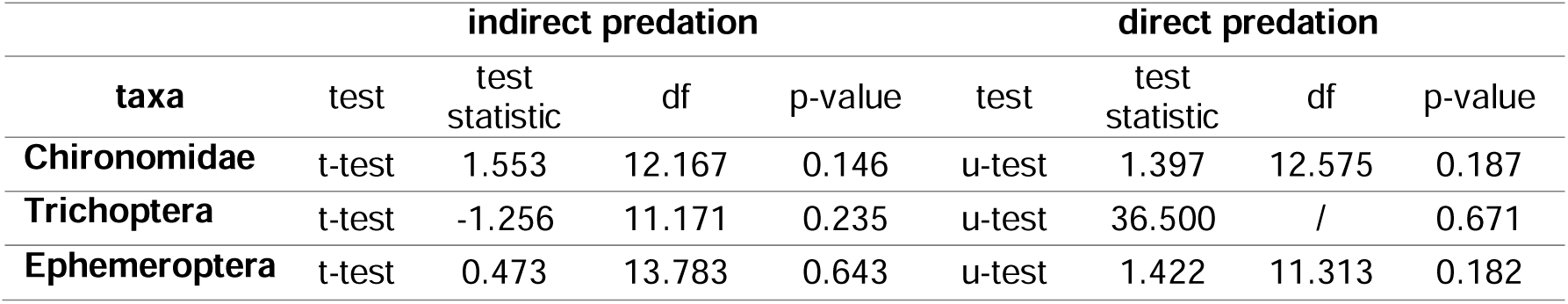
Influence of indirect and direct predation on mean body size during a nine-day mesocosm field experiment. Analysis used either a Student’s t-test for parametric data or a Wilcoxon rank-sum test (u-test) for non-parametric data. Predation was implemented via sticklebacks (*Gasterosteus aculeatus*) and bullheads (*Cottus rhenanus*).

| taxa | indirect predation |  |  |  | direct predation |  |  |  |
| --- | --- | --- | --- | --- | --- | --- | --- | --- |
|  | test | test statistic | df | p-value | test | test statistic | df | p-value |
| **Chironomidae** | t-test | 1.553 | 12.167 | 0.146 | u-test | 1.397 | 12.575 | 0.187 |
| **Trichoptera** | t-test | -1.256 | 11.171 | 0.235 | u-test | 36.500 | / | 0.671 |
| **Ephemeroptera** | t-test | 0.473 | 13.783 | 0.643 | u-test | 1.422 | 11.313 | 0.182 |

Larger Chironomidae (t = 1.397, df = 12.575, p = 0.187), Trichoptera (W = 36.5, p = 0.671), and Ephemeroptera larvae (t = 1.422, df = 11.313, p = 0.182) did not exhibit significantly increased drifting due to direct predation (Tab. 2).

### Predation influence on emergence number and size

We collected a total of 593 emergences in the net and drift samples. Out of these emergences, 566 individuals belonged to the Nematocera order. Among these, we identified 549 individuals, accounting for 92.58% of the total emergences, as Chironomidae imagos. The remaining emergences comprised 6 Simulidae (1.01%), 4 Chaoboridae (0.67%), 1 Psychodidae (0.17%), and 1 Staphylinidae (0.17%).

The number of emergences shows an initial increase in the first days, after which the frequency of emergence declines to only a few individuals that emerge on the eighth and ninth days (Fig. 7 A). In response to predation, the highest emergence numbers were observed on the third day. In contrast, under control conditions, the highest emergence numbers occurred on the fourth and sixth days (Fig. 7 A). Predation, whether direct or indirect, resulted in fewer invertebrates emerging overall (indirect predation: Chisq 4.264, df 1, Pr(>Chisq) 0.039, direct predation: Chisq 10.194, df 1, Pr(>Chisq) 0.001; Fig. 7 B). The size of the Nematocera, which represented the largest group among the caught emergences, was significantly smaller when the invertebrates were influenced by predation (t = 4.157, df = 12.942, p-value = 0.001, Fig. 7 C, D).

**Fig. 7:**
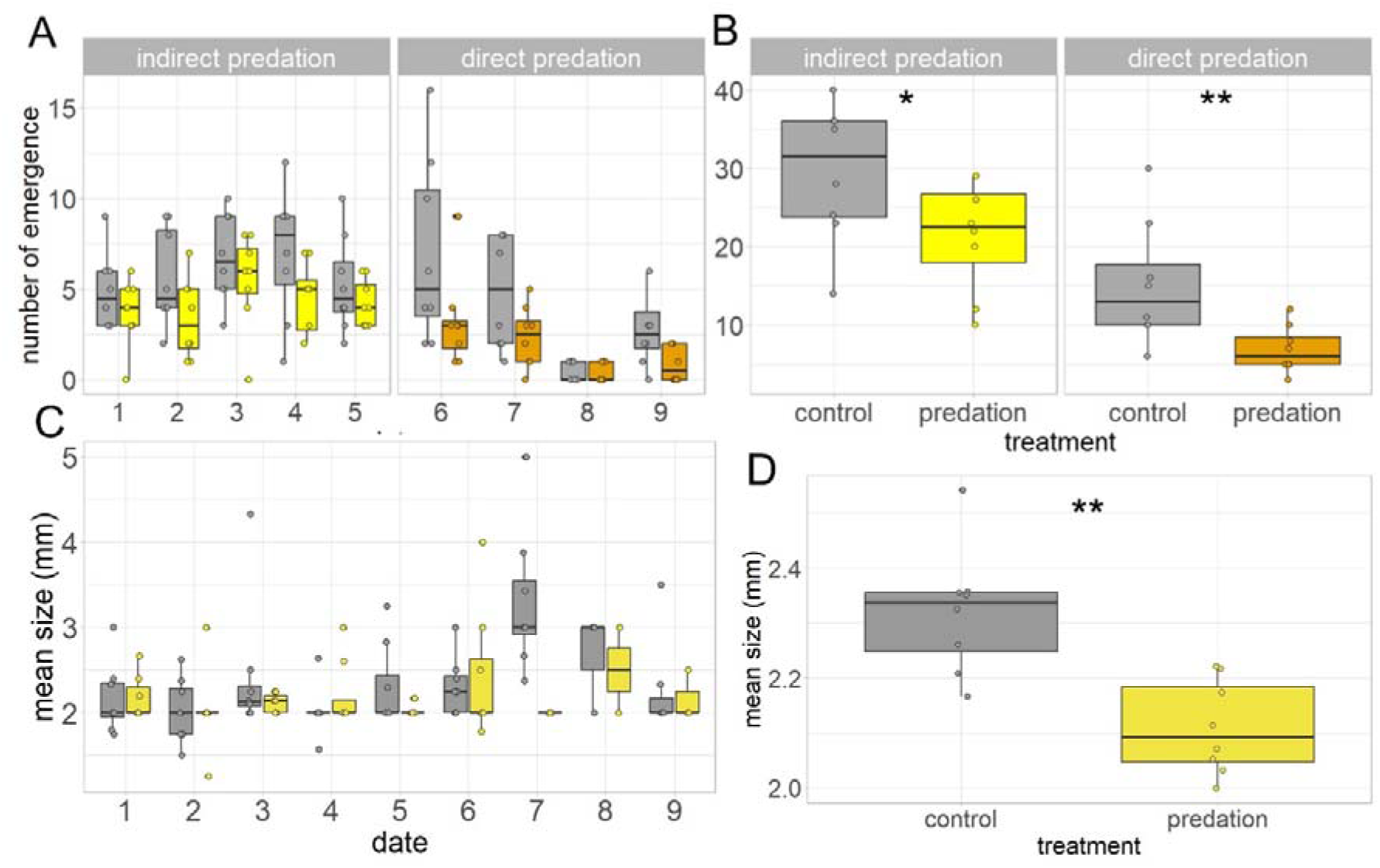
Influence of indirect (yellow) and direct predation (orange) on the number of emerging invertebrates in a field mesocosm experiment over nine days (A) and aggregated (B). Predation effects (dark yellow) on mean size (mm) of Nematocera over nine days (C) and aggregated (D). Predation was compared to control treatment without predation (grey). Water saturated with fish chemical cues from sticklebacks (*Gasterosteus aculeatus*) and bullheads (*Cottus rhenanus*) was added continuously starting from the first day of the experiment (yellow). At midnight of the fifth day, bullheads (*Cottus rhenanus*) were directly added as additional stimuli (orange). Indirect and direct predation significantly reduced emergence numbers (indirect predation: Chisq 4.264, df 1, Pr(>Chisq) 0.039, direct predation: Chisq 10.194, df 1, Pr(>Chisq) 0.001). The mean size of Nematocera is significantly reduced by predation (t = 4.157, df = 12.942, p-value = 0.001). Significant code: 0.1., 0.05 *, 0.01 **, < 0.001***.

## Discussion

### Drift response to indirect and direct predation

In our study, invertebrates responded to both indirect and direct predation. As the indirect predation treatment was fish-cue enriched water only, this points to the ability of invertebrates to detect predators through chemical cues (Weiss, 2019). Chironomidae larvae, Copepoda and Ostracoda reacted to indirect predation with increased drift, whereas Anomopoda, Nauplius larvae, *Gammarus* sp., Trichoptera, and Ephemeroptera larvae did not alter their drift behaviour. The lack of response in these taxa may be attributed to the natural fish-cue concentration in the Boye River, suggesting the potential for a predator-induced response in both treatments. However, some taxa did exhibit a response, indicating that our experimental setup likely intensified the fish cue signal. It has previously been shown, that increased cue levels, even against a background level of predation, can elicit antipredator responses (Brown et al., 2006). Alternatively, the taxa that did not respond to indirect predation by drifting could have also either reacted with an alternative antipredatory strategies such as reduced activity and hiding or they may not have perceived the chemical signal as a significant threat.

In comparison, direct predation was introduced after five days of continuous exposure to the chemical cues. Chironomidae larvae, *Gammarus* sp., Ostracoda, Nauplius and Trichoptera larvae reacted to direct predation with increased drift. While the preexposure could mean that the invertebrates’ sensitivity to these cues may have diminished (Brown et al., 2006), the direct placement of bullheads in the mesocosms most likely increased the fish-cue concentration. Moreover, direct predation involves not only chemical cues but also tactile and visual stimuli, as well as the potential for confrontation (Åbjörnsson et al., 1997). Therefore, we argue that direct predation should be perceived as a higher predation risk. This is supported by results of *Gammarus* sp., Nauplius and Trichoptera larvae showing increased drift only in response to direct predation.

However, this does not explain why the Copepoda did not significantly react if direct predation is a stronger stimulus. It is possible that most of the individuals had already left the system in response to indirect predation, reducing their numbers in the mesocosms and potentially decreasing the pool of individuals that would enter the drift. Insufficient numbers resulting in no measurable reaction towards predation may also explain the lack of response of Anomopoda and Ephemeroptera larvae. Ephemeroptera larvae, especially Baetidae, are typically known to increase their drift in response to predation, as observed in other studies (Culp et al., 1991; Peckarsky, 1996; Poff et al., 1991). Additionally, although we lack information about the number of prey directly consumed by predators during the direct predation phase, it is possible that the Copepoda were caught and eaten in the mesocosms starting from the sixth day, which could have impacted their numbers in the drift. This might also explain why the number of drifting Nauplius larvae dropped below that of the control treatment. However, the relevance of direct predation seems questionable, considering that bullheads are benthivores that usually do not prey on drifting organisms (Dahl, 1998a; Western, 1969). Furthermore, this should have more heavily impacted larger taxa, such as gammarids, due to the fish’s preferences (Western, 1969), rather than copepods, which constitute only a small percentage of their diet (Yoshiyama, 1980).

The distinct responses may also be explained by predation risk due to predator preferences for certain prey sizes and species. Depending on the predator’s size, gap width limitations, and hunting strategy, there may be a preference for larger prey (Holmes and McCormick, 2010; Persson et al., 1996; Slaughter and Jacobson, 2008), as larger prey typically offer greater mass and energy (Harper and Blake, 1988). Smaller taxa, such as Copepoda (< 1 mm), are commonly preyed upon by sticklebacks but not by the larger bullheads (Hynes, 1950; Western, 1969). Therefore, it may be conceivable that they do not respond to direct bullhead predation due to lower predation risk. Consequently, this could also explain why the average size of the drifting individuals was larger during direct predation. In contrast, except for the tendency observed in gammarid size, the sizes of most other taxa remained unaffected by predation. This could be due to the experimental design, sample processing or biological reasons. As the pump’s protection prevented animals over 5 mm in diameter from entering the system, this might have influenced the mesocosm population by allowing only smaller organisms to enter the system. We did seed organisms not filtered by size during the colonisation phase, however, the larger animals could have either left the system before the start of the stressor phase or could still be a percentual minority in comparison to the rest of the mesocosm population. The rounding of the size measurements may have also reduced differences that might otherwise have been statistically significant. It could also be that due to collecting drift samples every 24 hours rather than with a day vs night scheme, we missed any diurnal effects. Studies have shown that for example, the average size of drifting gammarids is larger at night (Andersson et al., 1986; Dahl, 1998a; Winkelmann et al., 2008), although that may depend on the predator. The nighttime drift helps reduce predation risk from visual hunting predators by decreasing the prey’s visibility.

Our findings suggest that predator avoidance behaviour in many species may be influenced by fish cues, with prey species being potentially able to sense changes in chemical cue concentration and assess predation risk (Botham et al., 2006; Brown et al., 2006; Kohler and McPeek, 1989; Lima and Dill, 1990; Peckarsky, 1996).

### Increased drift as the predator-induced response in all taxa

The predominant predator-induced response across the here investigated taxa was increased drifting. In many other studies, these same taxa showed contrasting behaviour, with reduced activity and less drifting (Bjærke et al., 2016; Gall and Brodie, 2009; Heuschele et al., 2020; Jourdan et al., 2016; Winkelmann et al., 2008). For example, although *Gammarus* sp. increased their drift under predation risk in our study, they are generally known to decrease activity and drift less (Andersson et al., 1986; Dahl, 1998a; Kasumyan, 2022). We anticipated at least some taxa to exhibit reduced drift, resulting in a “negative predator impact” as described by Sih and Wooster (1994). Nonetheless, the uniformity in responses observed in our study could be caused by different conditions, including the experimental setup, the habitat structure of the mesocosms, or the strength of our predator signal (Baumgärtner et al., 2003; Mathers et al., 2019).

The limited variety in behaviour may have, for example, been influenced by the scarcity of hiding spots within our mesocosms, primarily due to a sandy substrate and only one leaf litter bag. The substrate type and refuge opportunities have been demonstrated to significantly affect predator-induced behaviour (Baumgärtner et al., 2003; Penaluna et al., 2021). Moreover, the single leaf litter bag may not have provided an ideal refuge, given the confined space between the leaf litter inside the bag and its buoyancy.

In addition, the direct introduction of bullheads into the mesocosm may have intensified the predator signal beyond a critical threshold. This heightened threat perception could have prompted prey organisms such as the *Gammarus* sp. and Trichoptera larvae to resort to drastic measures such as fleeing the system via drift, rather than merely seeking refuge or reducing activity levels (Helfman, 1989). Drifting behaviour, which has been observed to occur under conditions of elevated predation risk and limited cover availability, may have been initiated as a survival strategy in response to these perceived threats (Penaluna et al., 2021) The behavioural responses of prey taxa could have also been shaped by the presence of multiple predators, each employing different hunting strategies and displaying preferences for particular prey species. These dynamics could have led to adaptive adjustments in prey behaviour, potentially manifesting as compromises or hierarchical predator-induced behaviours (McIntosh and Peckarsky, 1999).

### Short term predator-induced drift response

Our study revealed that increased predation pressure in riverine ecosystems elicits predator-induced drift behaviour in prey organisms. This behavioural response is initiated within the first 24 hours of exposure but diminishes considerably afterwards. We observed this decline in predator-induced response across several taxa, including Copepoda during indirect predation and Chironomidae larvae, *Gammarus* sp., and Nauplius larvae during direct predation. One reason for the relatively short-term response could be that the animals reacting to predation already fled the system within the first 24 hours, leaving no individuals to respond at a later point in time. This explanation may not fully hold, as evidenced by the response of Chironomidae to indirect predation, where their predator-induced drift response did not diminish. If most responsive Chironomidae larvae had already fled within the first 24 hours of the stimulus, we would not expect further response in the following days. Additionally, our system design allowed for the influx of new individuals from the river, suggesting that continuous responses should in general be possible. In the case of taxa with high upstream numbers and/or dispersal rates, previous studies have shown that populations are more likely to remain stable despite increased drift rates and mortality from predation pressure, thereby mitigating local predation effects (Cooper, 1990; Waters, 1972; Winkelmann et al., 2011).

### Emergence

In comparison, despite a high number of chironomids in the mesocosms (mean number of individuals: 152.75 ± 54.94), we observed only a few imagos emerging. This low emergence rate could be attributed high mortality rates during different live stages, as has been shown for various lotic insects (Huryn and Wallace, 2000; Otto, 1975; Parker and Voshell, 1979; Rutherford, 1986; Willis, and Hendricks, 1992).

In our study, emergence appeared to be time-dependent, with numbers peaking under control conditions around the fourth and sixth day before sharply declining, indicating synchronized emergence. Given their short adult lifespan, it is hypothesized that chemical communication among larvae signals the start of metamorphosis to synchronize emergence (Corbet, 1964). Without this synchronization, imagos would likely struggle to find mating partners in the limited time left in their adult stage. Depending on the species, imagos can live for only a few hours up to a few weeks. For instance, Ephemeroptera imagos live between a few hours up to a day or two, chironomid imagos typically live a few days, and Trichoptera imagos can survive up to several month before dying (Kriska, 2013). That we observed two peaks under control conditions could also be the result of protandry, the phenomenon that males of certain species emerge or mature earlier than females (Tokeshi and Reinhardt, 1996).

Predation exerted a noticeable influence on emergence, evidenced by a reduction in the number of emergences and their size. The decline in numbers could be attributed to a higher mortality rate among larvae and pupae due to predation pressure, or the deliberate prolongation of their larval stage avoiding predation (reviewed in Benard, 2004). By reducing their activity and foraging behaviour, the larvae increase their chances of staying undetected, but they also suffer from starvation (van Uitregt et al., 2012), which reduces growth and the necessary nutrients needed to safely initiate metamorphosis. Contrary to our expectation of a delayed metamorphosis, the peak in emergence number occurred earlier compared to the control treatment. However, previous studies have demonstrated that chironomids, which comprise a significant portion of our invertebrate population, tend to initiate metamorphosis earlier in the presence of predators as a strategy to escape unfavourable habitat conditions (Silberbush et al., 2019, 2015). This forced acceleration and earlier metamorphosis would also lead to reduced size (Stoks, 2001).

## Conclusion

In conclusion, both indirect and direct predation induced behavioural responses in most observed taxa, characterized primarily by increased drifting rates. This drift response may be attributed to the strength of the predation signal or conditions in our experimental setup, which provided few hiding opportunities. Size-dependent drift due to predation was also observed in the overall invertebrates, although on taxa level only gammarids were tendentially affected. The emergence of aquatic insects was similarly affected by predation, with reductions observed in both the number and size of emerging invertebrates. Our results show that chemical cues from fish have indirect effects on aquatic invertebrates potentially influencing dispersal and trophic interactions.

This alteration in prey dynamics may cascade through the food web, influencing the distribution and abundance of predators and prey and ultimately shaping the overall structure and function of the ecosystem.

## Acknowledgements

A sincere thank you is owed to Dr Christian Edler, Birgit Daniel (Bezirksregierung Düsseldorf, Dezernat 51) and Bernd Stemmer (Bezirksregierung Arnsberg, Fischereidezernat) for their invaluable assistance in catching the fish. We also thank Christoph D. Matthaei for his valuable advice. Special thanks go to the many people who actively supported us during the mesocosm experiment, including Alexander Rogalla, Theodore Domke, Antonia Domnik, Alexandra Hollstein, Johanna Knupfer, and Melina Rüngeler. We also wish to acknowledge and extend our sincere gratitude to Nicolai Bissantz for his assistance and expertise in addressing our statistical inquiries. This project is part of the Collaborative Research Centre 1439 RESIST (Multilevel Response to Stressor Increase and Decrease in Stream Ecosystems; www.sfb-resist.de) funded by the Deutsche Forschungsgemeinschaft (DFG, German Research Foundation; CRC 1439/1, project number: 426547801).

## Supplementary files

Supplementary tab. 1: Summary of glmmTMB (generalized linear mixed models using template model builder) results of drifting invertebrate numbers.

Supplementary tab. 2: Summary of glmmTMB (generalized linear mixed models using template model builder) results of drifting invertebrate numbers.

Supplementary tab. 3: Summary of glmmTMB (generalized linear mixed models using template model builder) results of drifting invertebrate numbers.

Supplementary tab. 4: Summary of the Anova results. Anova was applied to the results of the glmmTMB (generalized linear mixed models using template model builder) model results.

Supplementary tab. 5: Summary of the emmeans results.

Supplementary tab. 6: Summary of the glmmTMB (generalized linear mixed models using template model builder) and Anova results of emerging invertebrates.

## Statement of authorship

The project was planned by RT. The Initial planning for the implementation of the mesocosm experiment was done by FL, AJB and RT. They also organized the necessary administrative communication. Resources were organised by IMP. AMV, IMP, PMR implemented and improved the system. AMV investigated predator-specific behaviour. AMV, IMP, AJB, JM, MH and LW contributed to the statistical analysis of the data. AMV wrote the initial draft of the manuscript. All authors were involved in the refinement of the manuscript.

