## Supplementary tab. 1-6 for "Fish predation induces drifting and emergence in an experimental stream mesocosm system"

Supplementary tab. 1: Summary of glmmTMB (generalized linear mixed models using template model builder) results of drifting invertebrate numbers. Given are the indirect predation effects (p) affected by time (five days, day one to five). Predator-specific cues were introduced continuously into the system for indirect predator exposure (*Gasterosteus aculeatus* and *Cottus rhenanus*). To simulate direct predation, predator exposure was augmented by introducing bullheads (*Cottus rhenanus*) into the mesocosms. Bold font denotes tendential (< 0.05) and significant (< 0.1) p-values.

| taxa | coefficients | (Intercept) | predation | day2 | day3 | day4 | day5 | p:day2 | p:day3 | p:day4 | p:day5 |
| --- | --- | --- | --- | --- | --- | --- | --- | --- | --- | --- | --- |
| overall | Estimate | 90.039 | -31.471 | -33.402 | -52.354 | -49.878 | -55.747 | 16.223 | 26.115 | 29.519 | 26.028 |
|  | Std. Error | 6.562 | 8.180 | 7.598 | 6.819 | 6.929 | 6.733 | 9.329 | 8.509 | 8.718 | 8.364 |
|  | z value | 13.721 | -3.847 | -4.396 | -7.677 | -7.198 | -8.280 | 1.739 | 3.069 | 3.386 | 3.112 |
|  | Pr(>\|z\|) | **< 0.001** | **< 0.001** | **< 0.001** | **< 0.001** | **< 0.001** | **< 0.001** | ***0.082*** | **0.002** | **0.001** | **0.002** |
| Chironomidae | Estimate | 20.250 | -8.625 | -14.964 | -15.625 | -15.250 | -14.625 | 7.089 | 7.750 | 6.500 | **6.750** |
|  | Std. Error | 2.462 | 2.946 | 2.663 | 2.612 | 2.627 | 2.653 | 3.209 | 3.167 | 3.154 | 3.200 |
|  | z value | 8.226 | -2.928 | -5.620 | -5.982 | -5.805 | -5.514 | 2.209 | 2.447 | **2.061** | 2.109 |
|  | Pr(>\|z\|) | **< 0.001** | **0.003** | **< 0.001** | **< 0.001** | **< 0.001** | **< 0.001** | **0.027** | **0.014** | **0.039** | **0.035** |
| Gammarus sp. | Estimate | 10.547 | -3.782 | -0.939 | -2.889 | -1.587 | -4.770 | 0.792 | 2.233 | 3.119 | 3.116 |
|  | Std. Error | 1.391 | 1.842 | 1.603 | 1.464 | 1.533 | 1.344 | 2.046 | 1.917 | 2.016 | 1.758 |
|  | z value | 7.585 | -2.053 | -0.586 | -1.973 | -1.035 | -3.549 | 0.387 | 1.165 | 1.547 | 1.773 |
|  | Pr(>\|z\|) | **< 0.001** | **0.040** | 0.558 | **0.048** | 0.301 | **< 0.001** | 0.699 | 0.244 | 0.122 | **0.076** |
| Copepoda | Estimate | 30.400 | -12.100 | -13.699 | -21.985 | -19.808 | -19.850 | 7.761 | 11.448 | 12.559 | 9.892 |
|  | Std. Error | 2.852 | 3.501 | 3.380 | 2.985 | 3.100 | 3.096 | 4.154 | 3.713 | 3.900 | 3.823 |
|  | z value | 10.659 | -3.456 | -4.052 | -7.366 | -6.389 | -6.411 | 1.868 | 3.083 | 3.220 | 2.587 |
|  | Pr(>\|z\|) | **< 0.001** | **0.001** | **< 0.001** | **< 0.001** | **< 0.001** | **< 0.001** | ***0.062*** | **0.002** | **0.001** | **0.010** |
| Nauplius larvae | Estimate | 11.297 | -4.081 | -3.061 | -7.111 | -5.768 | -7.941 | 2.630 | 4.444 | 2.666 | 3.863 |
|  | Std. Error | 1.503 | 1.928 | 2.023 | 1.725 | 1.807 | 1.693 | 2.614 | 2.296 | 2.338 | 2.212 |
|  | z value | 7.514 | -2.116 | -1.513 | -4.122 | -3.192 | -4.690 | 1.006 | 1.936 | 1.140 | 1.746 |
|  | Pr(>\|z\|) | **< 0.001** | **0.034** | 0.130 | **< 0.001** | **0.001** | **< 0.001** | 0.314 | ***0.053*** | 0.254 | ***0.081*** |

Supplementary tab. 2: Summary of glmmTMB (generalized linear mixed models using template model builder) results of drifting invertebrate numbers. Given are the direct predation effects (p) affected by time (four days, day six to nine). Predator-specific cues were introduced continuously into the system for indirect predator exposure (*Gasterosteus aculeatus* and *Cottus rhenanus*). To simulate direct predation, predator exposure was augmented by introducing bullheads (*Cottus rhenanus*) into the mesocosms. Bold font denotes tendential (< 0.05) and significant (< 0.1) p-values.

| taxa | coefficients | (Intercept) | predation | day7 | day8 | day9 | p:day7 | p:day8 | p:day9 |
| --- | --- | --- | --- | --- | --- | --- | --- | --- | --- |
| overall | Estimate | 59.512 | -22.992 | -7.473 | -35.722 | -22.138 | 6.325 | 23.531 | 16.348 |
|  | Std. Error | 7.465 | 9.070 | 9.708 | 7.925 | 8.922 | 11.834 | 9.798 | 10.917 |
|  | z value | 7.972 | -2.535 | -0.770 | -4.508 | -2.481 | 0.534 | 2.402 | 1.498 |
|  | Pr(>\|z\|) | **< 0.001** | **0.011** | 0.442 | **< 0.001** | **0.013** | 0.593 | **0.016** | 0.134 |
| Chironomidae | Estimate | 9.169 | -4.807 | -2.511 | -5.873 | -7.245 | 4.624 | 3.573 | 5.802 |
|  | Std. Error | 1.494 | 1.805 | 1.969 | 1.722 | 1.653 | 2.540 | 2.117 | 2.103 |
|  | z value | 6.138 | -2.663 | -1.275 | -3.411 | -4.383 | 1.821 | 1.688 | 2.759 |
|  | Pr(>\|z\|) | **< 0.001** | **0.008** | 0.202 | **0.001** | **< 0.001** | ***0.069*** | ***0.091*** | **0.006** |
| Gammarus sp. | Estimate | 8.759 | -5.040 | -0.280 | -3.314 | -4.380 | -1.082 | 3.267 | 2.786 |
|  | Std. Error | 1.198 | 1.509 | 1.478 | 1.305 | 1.268 | 1.716 | 1.685 | 1.541 |
|  | z value | 7.311 | -3.339 | -0.189 | -2.540 | -3.455 | -0.630 | 1.939 | 1.808 |
|  | Pr(>\|z\|) | **< 0.001** | **0.001** | 0.850 | **0.011** | **0.001** | 0.528 | ***0.053*** | ***0.071*** |
| Copepoda | Estimate | 16.509 | -2.549 | -1.167 | -9.179 | -2.182 | -0.297 | 3.864 | -2.306 |
|  | Std. Error | 2.829 | 3.985 | 3.751 | 3.067 | 3.867 | 5.124 | 4.453 | 5.087 |
|  | z value | 5.836 | -0.640 | -0.311 | -2.993 | -0.564 | -0.058 | 0.868 | -0.453 |
|  | Pr(>\|z\|) | **< 0.001** | 0.522 | 0.756 | **0.003** | 0.573 | 0.954 | 0.385 | 0.650 |
| Nauplius larvae | Estimate | 7.552 | -1.859 | 1.335 | -4.347 | 3.658 | -2.171 | 5.112 | -2.140 |
|  | Std. Error | 1.396 | 1.821 | 1.978 | 1.497 | 2.309 | 2.439 | 2.157 | 2.828 |
|  | z value | 5.408 | -1.021 | 0.675 | -2.903 | 1.584 | -0.890 | 2.369 | -0.757 |
|  | Pr(>\|z\|) | **< 0.001** | 0.307 | 0.500 | **0.004** | 0.113 | 0.373 | **0.018** | 0.449 |

Supplementary tab. 3: Summary of glmmTMB (generalized linear mixed models using template model builder) results of drifting invertebrate numbers. Given are the indirect and direct predation effects. Due to low overall numbers and zero inflation, the data was aggregated, and the factor day was removed. Predator-specific cues were introduced continuously into the system for indirect predator exposure (*Gasterosteus aculeatus* and *Cottus rhenanus*). To simulate direct predation, predator exposure was augmented by introducing bullheads (*Cottus rhenanus*) into the mesocosms. Bold and italic font denotes tendential (< 0.05) and bold denotes significant (< 0.1) p-values.

|  | | indirect predation | | | | direct predation | | | |
| --- | --- | --- | --- | --- | --- | --- | --- | --- | --- |
| taxa | coefficients | Estimate | Std. Error | z value | Pr(>\|z\|) | Estimate | Std. Error | z value | Pr(>\|z\|) |
| Trichoptera | (Intercept) | 9.500 | 1.700 | 5.587 | **< 0.001** | 7.750 | 1.489 | 5.203 | **< 0.001** |
|  | predation | -1.125 | 2.294 | -0.490 | 0.624 | -3.250 | 1.789 | -1.816 | ***0.069*** |
| Ephemeroptera | (Intercept) | 8.625 | 1.095 | 7.873 | 0.000 | 7.000 | 1.193 | 5.869 | **< 0.001** |
|  | predation | -1.125 | 1.493 | -0.753 | 0.451 | -1.750 | 1.545 | -1.133 | 0.257 |
| Anomopoda | (Intercept) | 14.947 | 2.002 | 7.465 | **< 0.001** | 8.608 | 1.275 | 6.753 | **< 0.001** |
|  | predation | 1.855 | 2.905 | 0.639 | 0.523 | 2.909 | 1.951 | 1.491 | 0.136 |
| Ostracoda | (Intercept) | 10.625 | 1.633 | 6.507 | **< 0.001** | 7.875 | 1.500 | 5.251 | **< 0.001** |
|  | predation | -3.625 | 2.030 | -1.785 | ***0.074*** | -3.250 | 1.806 | -1.799 | ***0.072*** |

Supplementary tab. 4: Summary of the Anova results. Anova was applied to the results of the glmmTMB (generalized linear mixed models using template model builder) model results. Due to low overall numbers and zero inflation, the individual numbers of Trichoptera, Ephemeroptera, Anomopoda and Ostracoda were aggregated for indirect (day one to five) and direct predation (day six to nine), and the factor day was removed. Predator-specific cues were introduced continuously into the system for indirect predator exposure (*Gasterosteus aculeatus* and *Cottus rhenanus*). To simulate direct predation, predator exposure was augmented by introducing bullheads (*Cottus rhenanus*) into the mesocosms. Bold font denotes tendential (< 0.05) and significant (< 0.1) p-values.

|  | | indirect predation | | | direct predation | | |
| --- | --- | --- | --- | --- | --- | --- | --- |
| taxa | coefficients | Chisq | Df | Pr(>Chisq) | Chisq | Df | Pr(>Chisq) |
| overall | predation | 4.278 | 1 | **0.039** | 3.002 | 1 | ***0.083*** |
|  | day | 108.682 | 4 | **< 0.001** | 29.215 | 3 | **< 0.001** |
|  | predation:day | 14.283 | 4 | **0.006** | 7.750 | 3 | ***0.051*** |
| Chironomidae | predation | 10.285 | 1 | **0.001** | 1.240 | 1 | 0.265 |
|  | day | 56.429 | 4 | **< 0.001** | 28.328 | 3 | **< 0.001** |
|  | predation:day | 6.111 | 4 | 0.191 | 7.945 | 3 | **0.047** |
| Gammarus sp. | predation | 1.778 | 1 | 0.182 | 11.972 | 1 | **0.001** |
|  | day | 19.586 | 4 | **0.001** | 12.293 | 3 | **0.006** |
|  | predation:day | 4.442 | 4 | 0.349 | 10.533 | 3 | **0.015** |
| Copepoda | predation | 2.687 | 1 | 0.101 | 0.585 | 1 | 0.444 |
|  | day | 78.663 | 4 | **< 0.001** | 16.495 | 3 | **0.001** |
|  | predation:day | 12.322 | 4 | **0.015** | 3.136 | 3 | 0.371 |
| Nauplius larvae | predation | 1.681 | 1 | 0.195 | 0.323 | 1 | 0.570 |
|  | day | 36.149 | 4 | **< 0.001** | 13.962 | 3 | **0.003** |
|  | predation:day | 4.360 | 4 | 0.360 | 14.831 | 3 | **0.002** |
| Trichoptera | predation | 0.241 | 1 | 0.624 | 3.299 | 1 | ***0.069*** |
| Ephemeroptera | predation | 0.568 | 1 | 0.451 | 1.283 | 1 | 0.257 |
| Anomopoda | predation | 0.408 | 1 | 0.523 | 2.223 | 1 | 0.136 |
| Ostracoda | predation | 3.188 | 1 | ***0.074*** | 3.237 | 1 | ***0.072*** |

Supplementary tab. 5: Summary of the emmeans results. Emmeans was applied to the glmmTMB (generalized linear mixed models using template model builder) results when a tendentially or significant interaction was observed. Either indirect predation (indirect) or direct predation (direct) was compared with control over each day. Predator-specific cues were introduced continuously into the system for indirect predator exposure (*Gasterosteus aculeatus* and *Cottus rhenanus*). To simulate direct predation, predator exposure was augmented by introducing bullheads (*Cottus rhenanus*) into the mesocosms. Bold font denotes tendential (< 0.05) and significant (< 0.1) p-values.

| taxa | day | contrast | estimate | SE | df | z.ratio | p.value |
| --- | --- | --- | --- | --- | --- | --- | --- |
| overall | day 1 | no - indirect | 31.470 | 8.18 | Inf | 3.847 | **< 0.001** |
|  | day 2 |  | 15.250 | 6.43 | Inf | 2.373 | **0.018** |
|  | day 3 |  | 5.360 | 5.20 | Inf | 1.031 | 0.303 |
|  | day 4 |  | 1.950 | 5.49 | Inf | 0.355 | 0.723 |
|  | day 5 |  | 5.440 | 4.95 | Inf | 1.100 | 0.271 |
|  | day 6 | no - direct | 22.992 | 9.07 | Inf | 2.535 | **0.011** |
|  | day 7 |  | 16.667 | 8.27 | Inf | 2.016 | **0.044** |
|  | day 8 |  | -0.539 | 5.08 | Inf | -0.106 | 0.916 |
|  | day 9 |  | 6.643 | 6.92 | Inf | 0.960 | 0.337 |
| Chironomidae | day 6 | no - direct | 4.807 | 1.81 | Inf | 2.663 | **0.008** |
|  | day 7 |  | 0.183 | 1.79 | Inf | 0.102 | 0.919 |
|  | day 8 |  | 1.234 | 1.11 | Inf | 1.115 | 0.265 |
|  | day 9 |  | -0.995 | 1.08 | Inf | -0.922 | 0.356 |
| Gammarus sp. | day 6 | no - direct | 5.040 | 1.51 | Inf | 3.339 | **0.001** |
|  | day 7 |  | 6.120 | 1.45 | Inf | 4.224 | **<.0001** |
|  | day 8 |  | 1.770 | 1.35 | Inf | 1.317 | 0.188 |
|  | day 9 |  | 2.250 | 1.25 | Inf | 1.804 | ***0.071*** |
| Copepoda | day 1 | no - indirect | 12.100 | 3.50 | Inf | 3.456 | **0.001** |
|  | day 2 |  | 4.339 | 2.58 | Inf | 1.681 | ***0.093*** |
|  | day 3 |  | 0.652 | 1.83 | Inf | 0.356 | 0.722 |
|  | day 4 |  | -0.459 | 2.14 | Inf | -0.214 | 0.830 |
|  | day 5 |  | 2.207 | 2.01 | Inf | 1.099 | 0.272 |
| Nauplius larvae | day 6 | no - direct | 1.860 | 1.82 | Inf | 1.021 | 0.307 |
|  | day 7 |  | 4.030 | 1.88 | Inf | 2.149 | **0.032** |
|  | day 8 |  | -3.250 | 1.49 | Inf | -2.183 | **0.029** |
|  | day 9 |  | 4.000 | 2.37 | Inf | 1.684 | ***0.092*** |

Supplementary tab. 6: Summary of the glmmTMB (generalized linear mixed models using template model builder) and Anova results of emerging invertebrates. Anova was applied to the results of the glmmTMB model. Due to low overall numbers and zero inflation the individual numbers of Trichoptera, Ephemeroptera, Anomopoda and Ostracoda were aggregated for indirect (day one to five) and direct predation (day six to nine), and the factor day was removed. Predator-specific cues were introduced continuously into the system for indirect predator exposure (*Gasterosteus aculeatus* and *Cottus rhenanus*). To simulate direct predation, predator exposure was augmented by introducing bullheads (*Cottus rhenanus*) into the mesocosms. Bold font denotes tendential (< 0.05) and significant (< 0.1) p-values.

|  | glmmTMB | | | | | Anova | | |
| --- | --- | --- | --- | --- | --- | --- | --- | --- |
|  | coefficients | Estimate | Std. Error | z value | Pr(>\|z\|) | Chisq | Df | Pr(>Chisq) |
| indirect predation | (Intercept) | 29.477 | 2.991 | 9.856 | **< 0.001** |  |  |  |
|  | predation | -8.078 | 3.912 | -2.065 | **0.039** | 4.264 | 1 | **0.039** |
| direct predation | (Intercept) | 15.125 | 2.268 | 6.668 | **< 0.001** |  |  |  |
|  | predation | -8.250 | 2.584 | -3.193 | **0.001** | 10.194 | 1 | **0.001** |
